# Unraveling miRNA-Driven DNA Damage Response Networks in Pancreatic Adenocarcinoma: A Multi-Omics and Machine Learning Approach

**DOI:** 10.64898/2025.12.09.693279

**Authors:** Shayori Bose, Paramita Basak Upama, Sheikh Iqbal Ahamed

## Abstract

Due to the late detection, aggressive nature, and paucity of treatment options, pancreatic adenocarcinoma (PAAD) remains one of the most lethal cancers worldwide. MicroRNAs (miRNAs) are small non-coding RNAs that function as oncogenes or tumor suppressors. These are highly stable in the blood circulation, hence increasingly recognized as promising biomarkers for early cancer detection. We hypothesize that certain “driver miRNAs” regulate the expression of DNA Damage Response (DDR) genes. To test this, we have built a machine learning model to integrate multi-omics data from The Cancer Genome Atlas (TCGA) PAAD cohort, including copy number alterations, DNA methylation, gene expression, and microRNA (miRNA) expression, along with curated transcription factor interactions. We predicted around 2,000 miRNA-DDR gene interaction pairs; several of these were significantly enriched in a previously known experimentally validated miRNA-target interactions. Our analysis revealed several oncogenic and tumor-suppressive miRNAs that closely correlated with patient outcomes. Importantly, we were able to identify DDR genes that were highly targeted by multiple miRNAs and strongly correlated with patient overall survival. These findings enhance our understanding of the molecular mechanisms in PAAD and open new avenues for using miRNAs as disease biomarkers and suggest target genes for precision oncology.

## Introduction

Pancreatic adenocarcinoma (PAAD) is one of the deadliest cancers in the world, with a five-year survival rate of less than 12%^1^. The American Cancer Society estimates that in 2024, pancreatic cancer will be diagnosed in 66,440 patients in the United States, and 51,750 patients will die from the disease^1^. The prognosis of PAAD is considered dismal, with most patients presenting late in their disease course with local or distant metastatic disease^2,3^. Despite extensive genomic and transcriptomic characterization over the past decade, including analyses by The Cancer Genome Atlas (TCGA)^4^ and other large-scale^5,6^, the molecular drivers of PAAD progression and therapy resistance remain poorly understood. Standard of care treatments, most notably FOLFIRINOX and gemcitabine-based regimens, lead to modest survival benefits for metastatic pancreatic cancer patients, with a median overall survival of less than 12 months^7,8^. Therefore, the identification of robust biomarkers and regulatory networks is vital to enable early detection, prognostic stratification, and the design of targeted therapies.

MicroRNAs (miRNAs) are short, noncoding RNAs that function through complementary binding to target genes to typically repress the expression^9,10^. Acting as oncogenes or tumor suppressors, miRNAs influence diverse cellular processes, including proliferation, apoptosis, invasion, metastasis, and DNA repair^11,12^. Their stability in the circulation makes them easy to detect in various biological fluids, making them non-invasive biomarkers for cancer diagnosis, prognosis, and therapeutic monitoring^13,14^. Deregulated miRNA expression in pancreatic cancer is well documented, and several miRNAs (for example, miR-21, miR-155, miR-221, and the miR-200 family) have been identified as candidate prognostic^15,16^.

In PAAD, interactions between miRNAs and the DDR genes, which maintain genomic integrity, have long been speculated, but not well characterized.

The DNA damage response pathways represent a complex functions of genes that detect, signal, and repair various forms of DNA damage. This includes double-strand breaks, single-strand breaks, and replication fork stalling^17,18^. Dysregulation of DDR pathways is a hallmark of cancer progression and therapeutic resistance. For instance, deficiencies in homologous recombination repair DDR genes such as BRCA1, BRCA2, and PALB2 make them therapeutic targets of PARP inhibitors and platinum-based chemotherapy^19,20^. In pancreatic cancer, germline and somatic alterations in DDR genes occur in approximately 17-24% of patients with BRCA1 and BRCA2 mutations in 4-7% of cases^20,21^. Recent studies have shown that patients with germline BRCA and PALB2 mutations demonstrate favorable responses to platinum-based chemotherapy: one phase II trial showed response rates of 65-74% for gemcitabine plus cisplatin^22^. Additionally, studies showed that pancreatic cancer patients with BRCA1/2 mutations have Also, studies showed that pancreatic cancer patients with BRCA1 and BRCA2 mutations have a longer time to treatment discontinuation with FOLFIRINOX compared to gemcitabine plus nab-paclitaxel (9.3 vs 5.6 months, p = 0.028), suggesting potential predictive value of DDR alterations for platinum-containing therapies^23,24^. Finding DDR deficiency signatures as predictive biomarkers for FOLFIRINOX remains an area of active investigation.

We hypothesize that, given the established role of miRNAs in regulating DDR pathway components in other cancer types^25,26^, miRNA-mediated control of DDR genes could play a crucial role in PAAD biology, influencing both tumor development and therapeutic response.

Recent advances in multi-omics data generation technologies and computational models provide unprecedented opportunities to systematically map regulatory features of gene expression from different biological data modalities^27,28^. By integrating genomic, epigenomic, and transcriptomic data with curated protein-protein and miRNA-target interaction networks, it is now possible to uncover miRNA-driven regulatory circuits with functional relevance to cancer biology^29,30^. Machine learning approaches enhance the discovery of regulatory patterns that may be overlooked by traditional analytical methods^31,32^. These computational frameworks have proven particularly powerful in cancer research, enabling the identification of novel biomarkers, therapeutic targets, and drug resistance mechanisms across multiple cancer types^33,34^.

Previous experimental studies explored miRNA-DDR interactions in pancreatic cancer. For instance, miR-155 was found to target DNA repair genes, including MLH1 and MSH2, while miR-21 was found to modulate PTEN expression, affecting the PI3K/AKT pathway’s crosstalk with DDR signaling^35,36^. However, these studies have typically focused on individual miRNA-target pairs or small sets of interactions, lacking the comprehensive, systems-level approach necessary to fully characterize the miRNA-DDR regulatory landscape in PAAD. Furthermore, most existing studies have relied on cell line models or small patient cohorts, limiting the generalizability and clinical relevance of their findings^4,37^.

In this study, we sought to unravel the miRNA-DDR gene regulatory networks in PAAD using a comprehensive multi-omics framework combined with a machine learning approach. To achieve this, we developed a computational pipeline that integrates multi-omics data, including miRNA expression, gene expression, copy number alterations (CNA), and DNA methylation of 183 patients from TCGA. PAAD cohort, with curated transcription factors (TFs) and miRNAs interaction network. This pipeline applies a per-gene modeling strategy to apply machine learning to identify key “driver miRNAs” and their DDR gene targets (**Materials and Methods**).

Our integrative approach enables the systematic identification of miRNA-DDR regulatory circuits that are both statistically significant and biologically relevant. The resulting regulatory network highlights novel candidate biomarkers and regulatory hubs that not only deepen our understanding of PAAD pathogenesis but also provide a foundation for precision oncology strategies, including the development of miRNA-based therapeutics and the identification of patients most likely to benefit from DDR-targeted interventions such as PARP inhibitors and platinum-based chemotherapy^38,39^.

## Results

For each DDR gene, we built a predictive framework (**Figure 1**) combining gene expression, DNA methylation, CNA, TFs, miRNAs, and cis genes (genes within ±500 kb of the DDR locus). We ran the least absolute shrinkage and selection operator (LASSO) regression to predict each DDR gene’s expression using all candidate features. We then tune the regularization parameter (λ) with 10-fold cross-validation, selecting the simplest model with the lowest error. To stabilize feature selection, we ran LASSO 100 times per DDR gene and considered a miRNA to be a potential driver miRNA if it had a non-zero regression coefficient at least 70 times.

**Figure 1:**
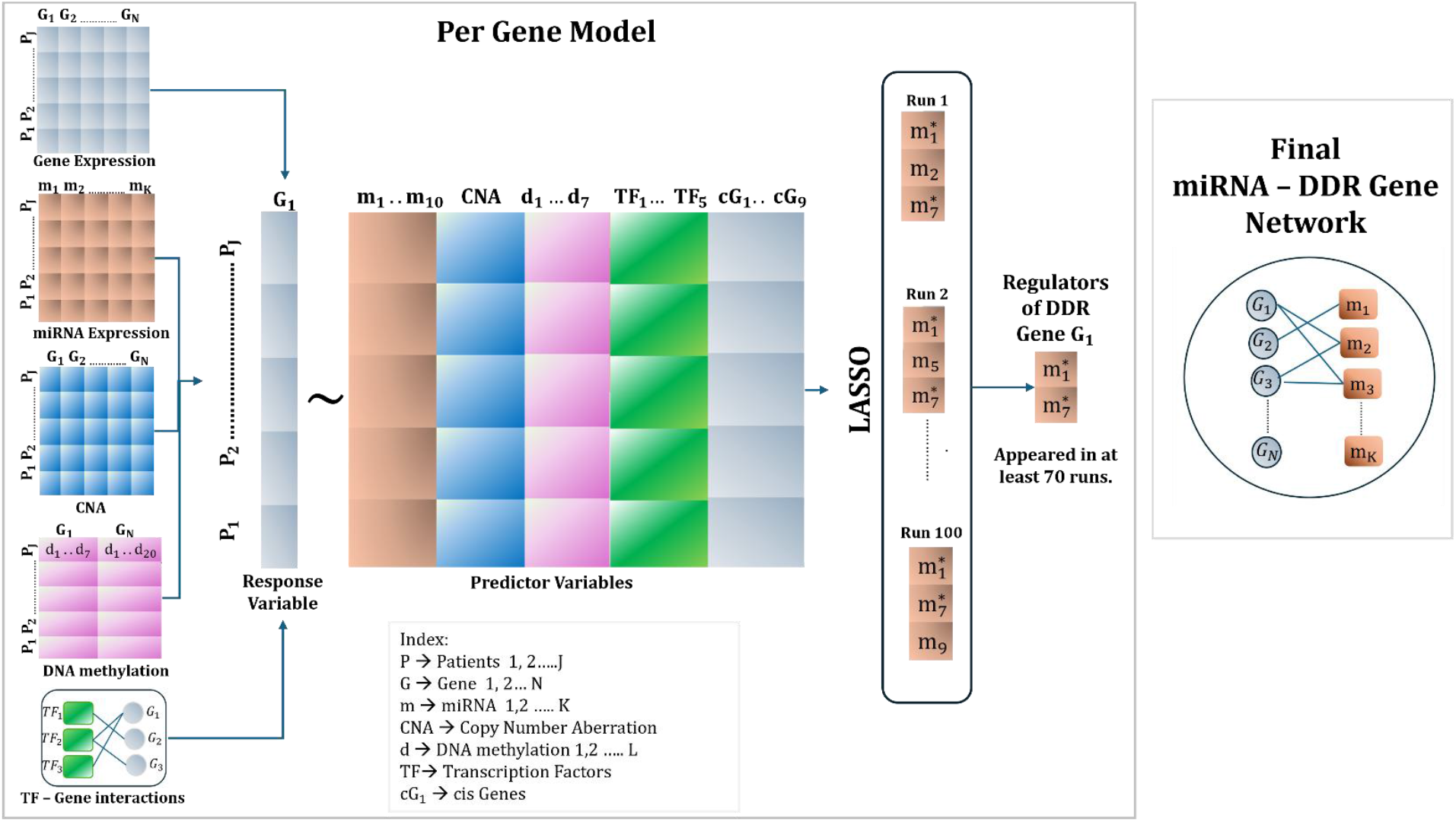
Machine learning pipeline integrating multi-omics datasets to predict miRNA-DDR gene network.

### Machine Learning pipeline identifies high-confidence miRNA-DDR gene regulatory networks

Our integrative multi-omics and machine learning pipeline yielded a high-confidence set of 2,203 miRNA-DDR gene interactions (814 miRNAs; 242 DDR genes). **Figure 2** depicts the miRNA distribution for each DDR gene. The hypergeometric test, performed for each selected miRNA targeting DDR genes, showed that these predicted interactions were significantly enriched (p-value < 0.05) in the reference set of experimentally validated miRNA-target gene pairs. Moreover, 20% of the identified miRNAs were known to target at least one DDR gene (p <0.05). **Table 2** shows specific miRNAs and the number of target DDR genes used in the hypergeometric test. The entire list can be accessed from **Supplemental Table 1**. Notably, several miRNAs were found to target several DDR genes exhibiting substantial regulatory breadth within DDR networks, such as mir-497 (targets 12 DDR genes, p = 0.04), mir-93 (targets 10 DDR genes, p = 0.01), and mir-375 (targets 6 DDR genes, p = 0.02). These results confirmed that the gene network reconstructed by our pipeline captured biologically meaningful interactions rather than random statistical associations, supporting its suitability for further analyses.

**Table 2:** A few miRNAs with the number of DDR genes and the p-values from the hypergeometric test.

| miRNA | No. of DDR genes | p-value |
| --- | --- | --- |
| mir-497 | 12 | 0.04 |
| mir-93 | 10 | 0.01 |
| mir-452 | 9 | 0.044 |
| mir-320z | 7 | 0.009 |
| mir-375 | 6 | 0.015 |
| mir-130b | 6 | 0.013 |
| mir-29a | 5 | 0.023 |
| mir-214 | 5 | 0.15 |
| mir-877 | 4 | 0.045 |
| mir-188 | 4 | 0.041 |
| mir-5659a | 3 | 0.033 |

**Figure 2:**
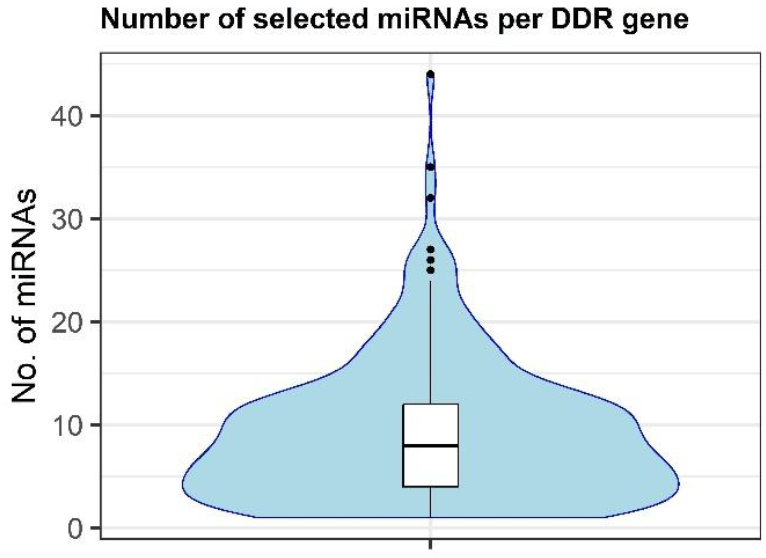
Violin plot showing the number of computed miRNAs per DDR gene.

### Identified miRNAs are significantly enriched for known oncogenic regulators

To test whether the miRNAs selected by our pipeline are known cancer-driving miRNAs, we checked their presence among oncogenic miRNAs annotated in oncomiRDB^40^. A two-sided Fisher’s exact test revealed highly significant enrichment (p = 3.3 × 10^−47^) in oncomiRDB, indicating that the inferred network is not only biologically plausible but also enriched for miRNAs with established roles in tumorigenesis.

This increases the confidence that novel miRNAs selected by our approach can be true drivers of PAAD progression.

### Selected miRNAs demonstrate strong prognostic value across multiple survival endpoints

Considering that DDR impairment is an integral part of tumorigenesis, we assessed whether selected miRNAs targeting DDR genes would be clinically relevant in patients. Adjusted Kaplan-Meier survival analyses were performed using inverse probability weighting (IPW) for age, gender, race, stage, and grade. Across four survival endpoints: Overall Survival (OS), Progression-Free Interval (PFI), Disease-Specific Survival (DSS), and Disease-Free Interval (DFI) (**see Materials & Methods**), a total of 143 miRNAs (18% of the total) were significantly associated with the prognosis of patients (adjusted log-rank p < 0.05). **Figure 3** shows representative Kaplan-Meier survival curves for the two significant miRNA biomarker candidates. For example, miR-375 expression clearly separated patients into high and low groups with significantly different OS (p = 0.031), while let-7b expression was associated with PFI (p = 0.001). **Table 2** presents the results of survival analyses for ten miRNAs that showed a significant association with the four endpoints. **Supplemental File 1** provides the survival plots for all 143 miRNAs significantly associated with at least one of the four clinical endpoints.

**Figure 3:**
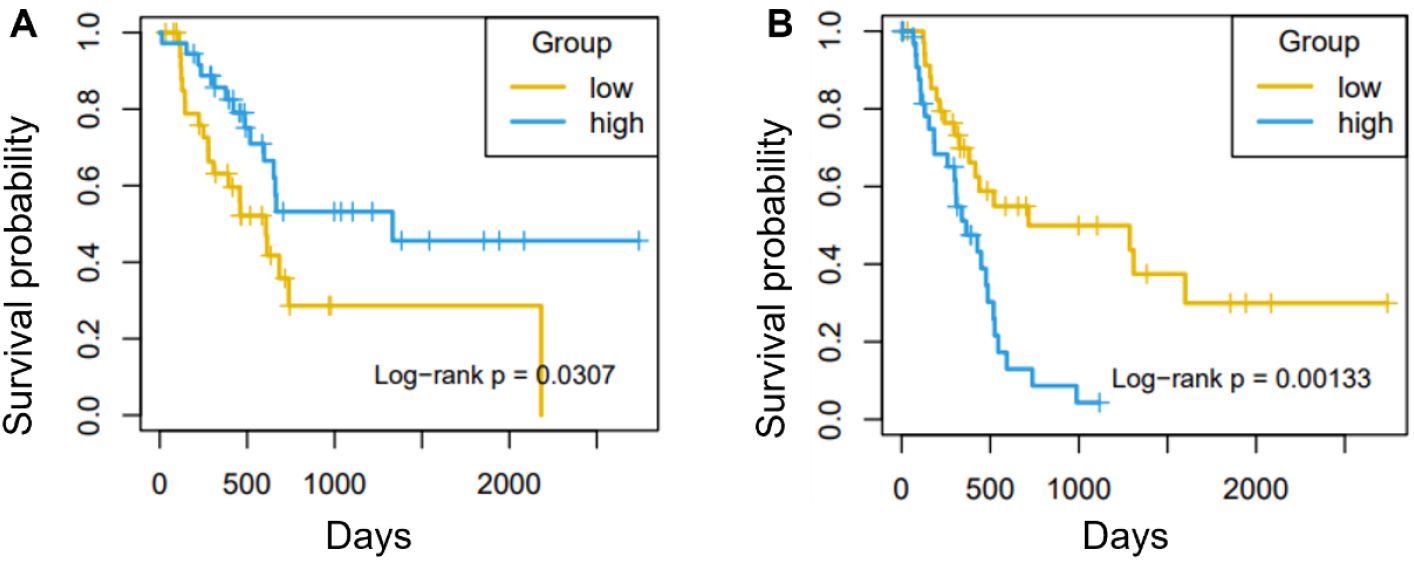
Kaplan-Meier plots for two high-degree miRNAs (i.e., miRNAs that target more than five DDR genes) in high-expression and low-expression patient groups in TCGA PAAD patients: A) miR-375 for OS and let-7b with PFI. Significant log-rank-test p-values showing the survival rate difference in the two groups.

**Table 2:** A few miRNAs with significant p-values showing the survival rate difference in the high and low expression groups for the four survival endpoints.

| miRNA | OS | PFI | DFI | DSS |
| --- | --- | --- | --- | --- |
| mir-375 | 0.03 | 0.01 | 0.01 | 0.03 |
| let-7b | 0.00 | 0.00 | 0.00 | 0.01 |
| mir-1179 | 0.02 | 0.01 | 0.02 | 0.02 |
| mir-1277 | 0.00 | 0.00 | 0.00 | 0.00 |
| mir-361 | 0.01 | 0.02 | 0.04 | 0.04 |
| mir-29b-1 | 0.00 | 0.00 | 0.00 | 0.00 |
| mir-1250 | 0.04 | 0.00 | 0.01 | 0.01 |
| mir-98 | 0.00 | 0.00 | 0.01 | 0.00 |
| mir-347b | 0.02 | 0.00 | 0.02 | 0.01 |
| mir-1301 | 0.00 | 0.00 | 0.04 | 0.03 |

These miRNAs play critical roles in pancreatic cancer progression and prognosis. miR-375 is frequently downregulated and functions as a tumor suppressor by targeting oncogenic pathways^41^. The let-7 family, including let-7b, suppresses KRAS signaling, and its loss is associated with poor differentiation and outcomes^42,43^. miR-1179, miR-361, miR-29b-1, miR-98, and miR-1301 have all been shown to inhibit proliferation and metastasis through targets such as TGFBR2, DUSP2/ERK, DNMT3B, MAP4K4, and RhoA^44–48^. Additionally, miR-1277 and miR-1250 have emerged from profiling studies as potential prognostic markers in pancreatic cancer, though their functions require further validation^49,50^. Together, these miRNAs encompass known tumor suppressors and novel potential biomarkers in pancreatic cancer. This further supports the use of DDR-related miRNAs as potential biomarkers for patient stratification and treatment planning.

### Network analysis identifies high-degree DDR gene hubs and shows their regulatory potential

Analysis of the miRNA-DDR gene interaction network demonstrated that the miRNAs did not target the DDR genes uniformly. Some DDR genes represented hubs with a high degree as they were targeted by multiple miRNAs. **Figure 4** shows that POLD3 and RIF1 gene targets, which are regulated by 15 and 35 miRNAs. **Supplemental Table 2** shows the list of forty high-degree DDR genes (≥ 15 targeting miRNAs). These hubs may represent vulnerable genes in the DDR pathways that, when regulated, drive genomic instability and tumor progression, making them attractive candidates for therapeutic intervention.

**Figure 4:**
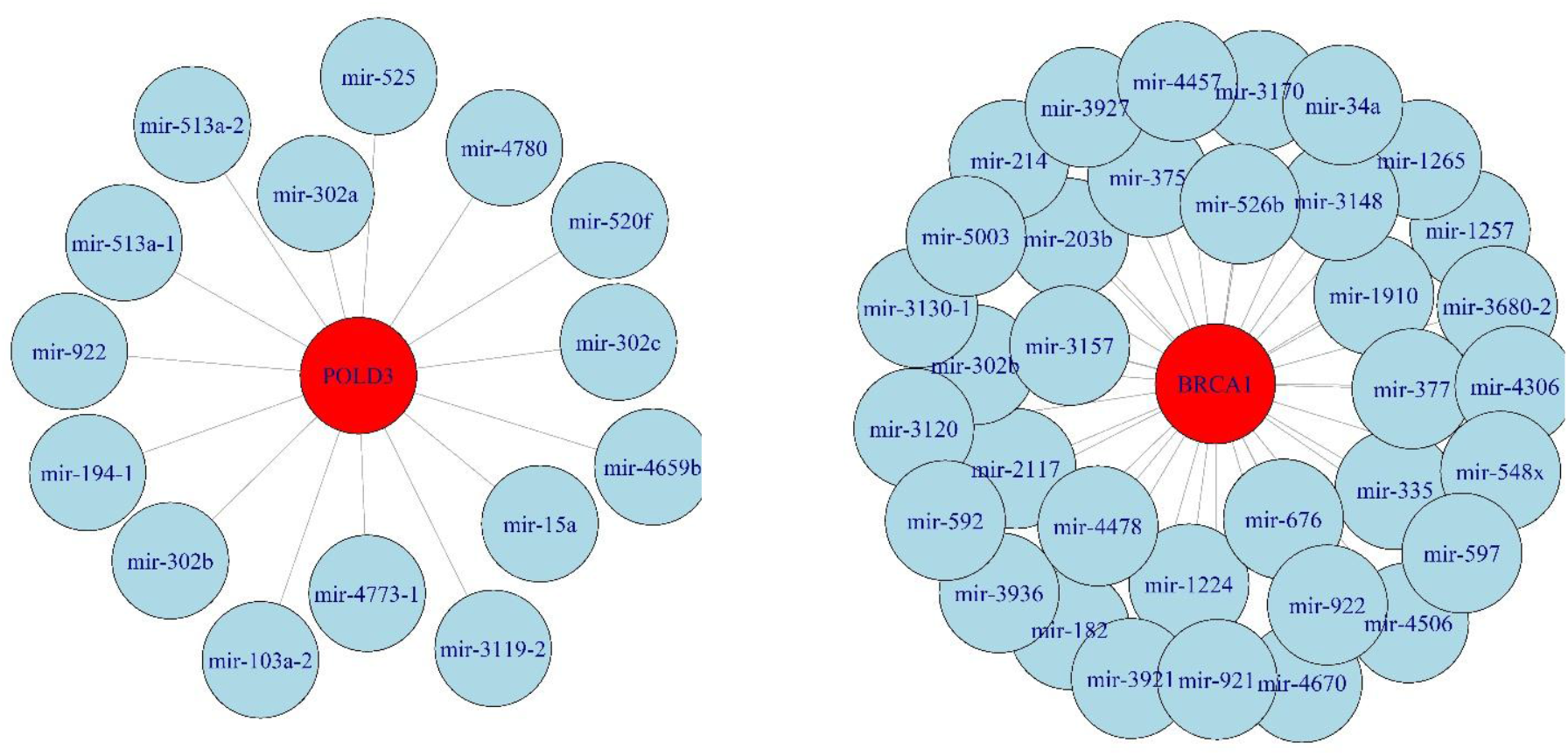
miRNA-DDR interaction network for POLD3 and RIF1.

### Functional enrichment confirms that the high-degree DDR genes are central in core DNA repair pathways

To elucidate the biological nature of genes that are targeted by at least five miRNAs, we performed functional enrichment analyses. KEGG^42^ pathway enrichment analysis also showed a strong overrepresentation of the canonical DNA repair and genome maintenance pathways, such as Fanconi anemia, base excision repair, nucleotide excision repair, homologous recombination, mismatch repair, cell cycle and DNA replication (**Figure 5A**). Gene Ontology (GO)^51^ enrichment further corroborated these results, showing significant enrichment of the following biological processes: double-strand break repair, DNA recombination, regulation of cell cycle phase transition, response to radiation and DNA-templated replication (**Figure 5B**). These results confirm that the highly targeted DDR genes identified by our pipeline are interconnected within fundamental pathways for genomic integrity, supporting their potential impact on PAAD.

**Figure 5:**
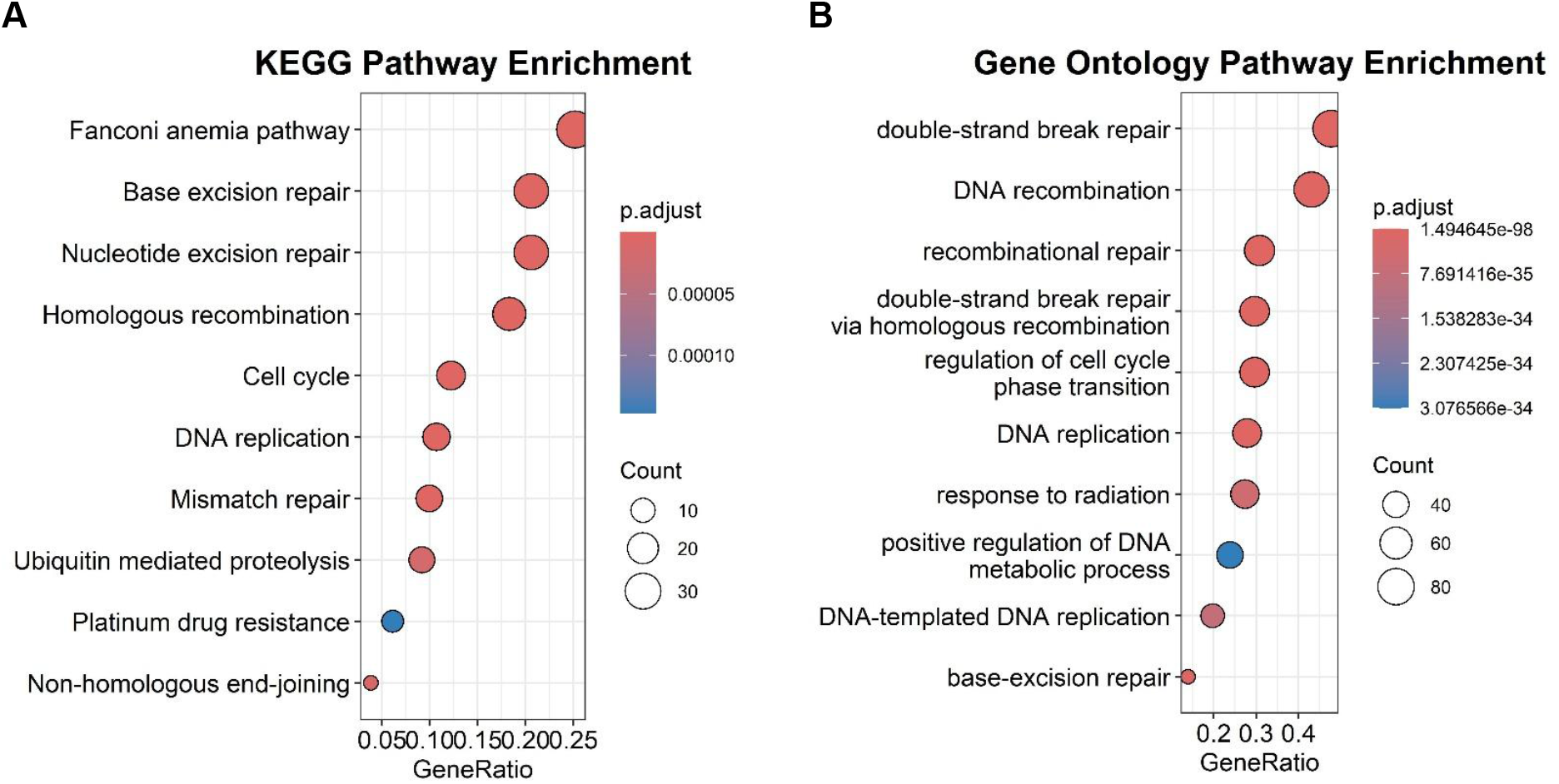
Enriched (adjusted p-value < 0.05) A) KEGG pathways and B) GO-based biological processes pathways for the DDR genes targeted by ≥ 5 miRNAs.

Our multi-omics and machine-learning-based framework described here has revealed the landscape of miRNA-DDR gene interactions, enriching for biologically valid, clinically relevant, and prognostically informative interactions. Specifically, the framework enriched for experimentally validated miRNA-target pairs captured substantial overlaps with known oncogenic miRNAs, and revealed strong prognostic associations across multiple survival endpoints for both miRNAs and their DDR gene targets. It also revealed the existence of high-degree regulatory hubs in which miRNAs appeared to interact with large numbers of genes, forming central nodes in core DNA repair pathways. These findings broaden our understanding of DDR regulation in PAAD, and nominate candidate biomarkers and regulatory nodes for further experimental validation and potential therapeutic intervention.

## Discussion

In this study, we designed a computational framework to identify miRNA-DDR gene networks in PAAD. By integrating miRNA expression, gene expression, DNA methylation, and copy number data with curated regulatory interactions, we identified biologically validated miRNA-DDR gene interactions and delineated hubs of DDR genes under the control of multiple miRNAs. Importantly, several miRNAs were prognostically relevant for multiple clinical endpoints of the disease.

Selected miRNAs such as miR-497, miR-93, and miR-375 have previously been identified in tumorigenesis, which gives strong biological support to our findings. POLD3, RIF1, and other DDR-related genes targeted by multiple miRNAs are potential high-degree hubs for replication stress prevention and genome maintenance. Prognostic analyses (e.g., let-7b, miR-375) further revealed that miRNAs could serve as biomarkers and therapeutic targets. The DDR-related hallmark of PAAD progression can be attributed to the miRNA-driven regulation pathways.

While there are clear strengths to our approach, our analysis of bulk tumor data from a moderately sized cohort is limited in resolution in terms of the intratumoral heterogeneity and microenvironmental influences. Recent advances in single-cell RNA-seq and single-cell ATAC-seq offer the opportunity to delineate cell-type-specific miRNA-DDR regulatory programs, particularly as they distinguish between malignant epithelial cells and stromal and immune compartments. At the same time, larger multi-omics clinical cohorts, capturing key transcriptomics, proteomics, chromatin accessibility, and spatial profiling information, will be needed to confirm and refine these regulatory signatures. Such integrative approaches will reveal how miRNA regulation intersects with transcriptional, epigenetic, and translational regulation to control the activity of DDR genes and promote or impair therapeutic responsiveness.

Together, our findings broaden the scope of post-transcriptional regulation of DDR pathways in pancreatic cancer and nominate key miRNAs and DDR gene hubs with prognostic and therapeutic implications. Future studies using single-cell resolution, expanded multi-omics data sets, and functional perturbation experiments will be needed to define the mechanistic basis of these networks and translate them into actionable biomarkers and therapeutics.

## Materials and Methods

### Download DDR gene list

We curated a DDR gene list from Knijnenburg et al.^52^, which consists of 274 genes from MSigDB v5.0 and other expert-curated resources. Of these, 211 (76%) span nine major DDR pathways: base excision repair, nucleotide excision repair, mismatch repair, the Fanconi anemia pathway, homology-dependent recombination, non-homologous end joining, direct damage reversal/repair, translesion DNA synthesis, and nucleotide pool maintenance.

### Download and pre-process multi-omics data

Genomic and transcriptomic data for PAAD were retrieved from the TCGA database using the *TCGABiolinks*^53^ R/Bioconductor package. The study cohort consisted of 183 patients. mRNA expression data were downloaded as raw counts from Illumina HiSeq and subsequently converted to RPKM values after filtering out lowly expressed genes. miRNA expression quantification files (hg19.mirbase20.mirna) and isoform files (hg19.mirbase20.isoform) were retrieved with ≥0.01 RPM in at least 30% of the samples. We harmonized the miRNA expression data with TCGA by mapping the identifiers to MIMAT IDs and by removing redundant 3p/5p suffixes from mature miRNA names. Masked somatic copy number variation (CNV) profiles from the Affymetrix SNP Array 6.0 platform were converted to gene-centric CNV values aligned to hg38 using the *CNTools*^54^ R package. DNA methylation data for PAAD samples were sourced from the Illumina HumanMethylation450 platform. Probes missing all sample values were excluded. SNP information from dbSNP was added using the *minfi* R package, and probes containing SNPs at the CpG interrogation site or at single-nucleotide extension were removed. The remaining probes were then filtered to retain only those that map to the following gene regions: gene body, first exon, TSS1500, TSS200, and 5’ UTR. Filtered probes were mapped to the Ensembl gene IDs using the Biomart annotation data, thereby producing gene-specific methylation probe sets for downstream analyses.

### Integrate miRNA-target and TF-target Interactions

We retrieved computationally predicted miRNA-target interactions from DIANA-microT^55^, ElMMo^56^, miRDB^57^, PicTar^58^, and TargetScan^59^. For TargetScan, we applied a conservation filter and retained only the top 20% of binding sites. These miRNA names were aligned to TCGA MIMAT identifiers to resolve naming and strand-specific differences. For each DDR gene, a gene-specific miRNA expression matrix was constructed that included all expressed putative miRNAs. A total of 274 DDR genes had at least one miRNA in the target dataset, corresponding to 1,704 unique miRNAs.

We obtained high-confidence TF–target interactions from DoRothEA^60^, a resource that integrates various lines of evidence, including literature curation, ChIP-seq data from the ENCODE^61^ project and others. TF and target HGNC symbols were linked to Ensembl gene IDs based on Biomart annotations; only TFs with detectable expression in PAAD RNASeq samples were retained. For each DDR gene, a TF expression matrix was created and stored for integration with other molecular predictors. Across the PAAD cohort, a total of 1,512 TFs regulating 28,214 target genes were retained.

### Per-DDR gene machine learning model

For each DDR gene, we developed an integrative model that considered the combined effects of mult-omics datasets, including gene expression, DNA methylation, CNA, miRNAs, and cis-genes (defined as genes located within ±500 kb of a given DDR gene according to GENCODE annotations) (Figure 1) and a TF-target interaction network. We included DNA methylation probes and CNA segments within 500 kb of the locus as candidate predictors. In addition, only TFs and miRNAs expressed in the PAAD cohort were retained, and miRNA names were harmonized to TCGA MIMAT identifiers.

The model for each DDR gene was built by merging all candidate predictor matrices by sample ID, removing empty rows and columns, and keeping only those samples that had complete data in all predictor types. Gene expression, DNA methylation, CNA, TF expression, cis gene expression, and miRNA expression values were treated as independent variables to predict DDR gene expression (response variable). The feature sets for each gene thereby encompassed both cis and trans information on DDR regulation.

We implemented a least absolute shrinkage and selection operator (LASSO) regression^62^ to model DDR gene expression as a function of all candidate features. Optimal λ was determined as the most parsimonious model within one standard error of minimal error via 10-fold cross-validation. To address the possibility of instability, we repeated the LASSO procedure 100 times for each DDR gene. MiRNAs with non-zero coefficients in at least 70 out of 100 runs were considered potential regulators, which is consistent with the previous studies using the LASSO method^63,64^.

### Survival analyses

To determine the clinical relevance of identified miRNAs, we performed survival analyses across four endpoints defined in previous literature. For OS, patients who had died from any cause were considered as having experienced the event, while all other patients were censored; PFI considered patients as having an event if they experienced a new tumor occurrence, including disease progression, local recurrence, distant metastasis, new primary tumor, or death with cancer in the absence of a new tumor event, as well as cases where the type of new tumor event was not specified. All other patients were censored. DFI followed the same definition as PFI but additionally included patients censored due to new primary tumors in other organs; patients who died with tumors without a new tumor event and those with stage IV disease were excluded; for DSS, the longest available time among disease-specific survival days, last contact days, or death days was used to determine whether the event occurred, with all remaining patients considered censored^65^. Survival analyses were performed using the Kaplan-Meier method with the *survival* R package, and visualized with *survminer*. MiRNAs or DDR genes were considered prognostic if the log-rank test^66^ produced a p-value < 0.05.

### Enrichment of cancer-related miRNAs

To evaluate the oncogenic relevance of identified miRNAs, we compared our selected set to a curated list of cancer-associated miRNAs from the oncomiRDB^40^ database, harmonized with miRBase^67^ nomenclature. A two-sided Fisher’s exact test established enrichment, confirming the biological significance of the miRNAs identified by our modeling.

## Code and Data Availability

The full source code for the machine learning model, including scripts for figure and table generation, model evaluation, and the supplemental materials, can be accessed from https://github.com/shayorib/Pancreatic_Cancer. Pancreatic cancer molecular and clinical data were obtained from TCGA via the Genomic Data Commons (GDC) Data Portal: https://portal.gdc.cancer.gov/.

